# A Causal Role of the Right Dorsolateral Prefrontal Cortex in Random Exploration

**DOI:** 10.1101/2024.04.28.591514

**Authors:** Armin Toghi, Mojtaba Chizari, Reza Khosrowabadi

**Affiliations:** Institute for Cognitive and Brain Sciences, Shahid Beheshti University, Tehran, Iran

## Abstract

Our brain faces the dilemma of exploiting familiar options to gain immediate rewards or exploring new options to increase probable future rewards, in many real-life decisions, and solves it by direct and random exploration. Previous studies show that these two explorative strategies have dissociable neural correlates in the brain. Using the continuous theta burst stimulation (cTBS) and horizon task, we investigate the causal role of the right dorsolateral prefrontal cortex (rDLPFC) in direct and random exploration. Twenty-five healthy right-handed adult participants underwent cTBS, and vertex stimulation sessions, and then completed the horizon tasks. Both model-free and model-based analysis showed that cTBS over rDLPFC selectively reduced random exploration, but not direct exploration. This suggests a causal role for rDLPFC especially in random exploration, and further supports dissociable neural implementations for direct and random exploration. © 2024 The Author(s)

## 1. Introduction

In almost every decision we make in everyday life, we are choosing between two categories of options: familiar options that you know their values, and the novel, which has uncertain outcomes. A familiar option like choosing our favorite food has a high immediate reward compared to choosing a new food we have never tested before. However, adaptive behavior requires exploring new options to increase probable future reward [Cohen et al., 2007]. Thus, in a dynamic environment like the real world there has to be a balance between choosing a familiar option with a known expected reward (Exploitation) and choosing new options to test their probable reward (Exploration). This balance, commonly referred to as the exploration-exploitation tradeoff, constitutes a fundamental computational challenge that the brain faces in everyday life [Cohen et al., 2007, Wilson et al., 2014].

Previous studies have identified two primary approaches that humans utilize to address the exploration-exploitation tradeoff. These approaches can be classified into two groups: direct and random exploration [Wilson et al., 2014]. Direct exploration involves choosing an option that is more informative about underlying values [Auer et al., 2002]. Meanwhile, random exploration occurs when there is stochasticity in decisions [Thompson, 1933], or when there is complete uncertainty regarding the available options [Gershman, 2018, Tomov et al., 2020], potentially leading an agent to select a suboptimal choice.

In recent years, there has been converging evidence showing dissociable neural implementation underlying these two explorative strategies in the human brain. The first line of research comes from studies supporting the differential role of dopamine and noradrenaline in mediating exploration [Chakroun et al., 2020, Cremer et al., 2023, Dubois et al., 2021, Fan et al., 2023]. Pharmacological intervention (manipulation) in human participants supports the hypothesis that dopamine mediates direct exploration [Chakroun et al., 2020, Cremer et al., 2023], while norepinephrine controls random exploration [Dubois et al., 2021, Fan et al., 2023]. The second line of research comes from studies that showed the dissociable role of the right frontopolar cortex (rFPC) and dorsolateral prefrontal cortex (rDLPFC) in controlling direct and random exploration, respectively. Previous neuroimaging studies showed that rFPC drives information-seeking and direct exploration [Badre et al., 2012, Hogeveen et al., 2022a, Mansouri et al., 2017, Tomov et al., 2020], while rDLPFC drives random exploration [Jahn et al., 2023, Tomov et al., 2020]. Moreover, transcranial magnetic stimulation (TMS) over the rFPC decreases direct exploration while intact random exploration [Zajkowski et al., 2017b]. However, unlike the rFPC, there is no direct causal evidence for the involvement of the rDLPFC exclusively for random exploration.

The right DLPFC has been linked to various underlying functions that facilitate exploratory behaviors, such as accumulation of uncertainty [Huettel et al., 2005], or encode the immediate and future value of choice options [Hogeveen et al., 2022b, Tang and Averbeck, 2022], before making a decision. Moreover, right DLPFC is associated with other related constructs like risk-taking and impulsivity [Cho et al., 2010, 2012, Dantas et al., 2023, Obeso et al., 2021]. However, brain stimulation techniques over right DLPFC yielded mixed results in decision-making tasks [Ngetich et al., 2020]. While some studies showed that inhibiting rDLPFC decreases impulsivity [Cho et al., 2010], others support increasing risk-taking behavior [Dantas et al., 2023, Knoch et al., 2006]. Although these definitions (e.g., risk-taking and impulsivity) are related to explore-exploit behavior, studies have not converged to the operational definitions of exploration-exploitation behavior [Wilson et al., 2021].

Furthermore, identifying direct and random exploration in exploration tasks remains challenging due to various psychological confounds, such as the reward-information confound, and individual differences in ambiguity preference [Wilson et al., 2014]. Horizon tasks are one of the perfectly designed tasks for distinguishing between direct and random exploration, which solve the problem of measuring direct exploration and assessing the contribution of choice variability to random exploration [Wilson et al., 2014].

Here, we used continuous theta-burst stimulation to selectively inhibit the rDLPFC in participants performing horizon tasks. While this task selectively differentiates between direct and random exploration, we could directly investigate the role of the right DLPFC in direct and random exploration.

We find that downregulating the excitability of rDLPFC significantly decreases the random exploration (i.e., probability of choosing options with low mean). At the same time, direct exploration didn’t change as measured by both model-free and model-based analysis. Together, these results support the role of rDLPFC in decreasing the sensitivity to average rewards in total uncertainty conditions and driving random exploration.

## 2. Experimental Procedures

### 2.1. Participants

For estimating the sample size, a priori power analysis was conducted using G*Power version 3.1.9.7 [Faul et al., 2007]. Based on an exploratory study of the non-invasive brain stimulation meta-analyses, we put 0.62 for power [Mitra et al., 2019]. With a significance criterion of α = .05 and effect size = 0.4 (considered to be large using Cohen’s (1988) criteria), the minimum sample size needed with this effect size is N = 20 for repeated measure ANOVA. Thus, we recruited 30 healthy right-handed, adult volunteers (14 female, 18-32) for this study. Participants were recruited from paper flyers distributed at the Shahid Beheshti University and the university campus.

None of the subjects had a history of brain injury, head trauma, psychiatric or neurological disorders, and had no previous experience with horizon task, or TMS, with normal or corrected-to-normal vision. Subjects were screened for depression and anxiety with the use of the Depression Anxiety Stress Scales – DASS-21 with an exclusion criterion of the score of 10 for each subscale. Furthermore, we don’t include economic and technical science students with prior knowledge of decision theories, which might induce bias in the study.

Three participants were excluded due to discomfort during stimulation (2 female, 1 male). Two participants were excluded due to high resting motor threshold (2 female, 0 male), and two participants were excluded due to chance level performance in experimental sessions. The final data contains 23 participants (11 female, mean age: 23.84, sd: 2.76) with completed data.

All participants were informed about the potential risks of TMS and signed informed consent before the first session. The study was conducted under the Helsinki regulations and the ethics of the experiment was approved by the university and the Humanities Ethics Committee with reference number IR.SBU.REC.1402.188.

### 2.2. TMS protocol

We used continuous theta burst stimulation protocol (cTBS) with Magstim Pro 2 devices, which contain a continuous 40-sec train of 600 pulses, with short bursts (3 stimuli) of 50 Hz pulses repeated at theta range (5 Hz). This protocol is expected to decrease cortical excitability for up to 50 minutes [Wischnewski and Schutter, 2015].

Our protocol contains two sessions (Figure 2A). In the first session, participants got familiar with the task, and then we obtained a resting motor threshold, defined as the lowest stimulation intensity needed to elicit a visible contraction of the abductor pollicis brevis (APB) in five out of ten pulses after stimulating the motor cortex. Then stimulation was applied based on individual resting motor threshold. While previous studies mostly applied an 80 % individual resting motor threshold, recent studies suggested higher doses for proper stimulation conditions. That’s why, we applied a 100 % individual resting motor threshold [Hogeveen et al., 2022a]. To find stimulation sites, we used the 10-20 EEG international system. One session was the vertex, where we put the TMS coil on the Cz position, and for stimulating the rDLPFC, the coil was located in the F4 position (Figure 2B). Although using neuronavigation methods with MRI coordinates is better for localizing the specific brain area, DLPFC is a large structure and we considered its localization with a general 10–20 EEG system to be sufficient to reduce cortical excitability.

### 2.3. Procedure

Each participant undergoes two TMS experimental sessions with an interval of at least 5 days. In the first session, they went through 10 training games to get familiar with the task. Next, we obtained a resting motor threshold and then the stimulation took place. The order of stimulation was counterbalanced across participants. Participants started the Horizon task 5 minutes after stimulation, to allow for the downregulating effects of cTBS to take place.

### 2.4. Horizon Task

The horizon task was designed based on Wilson and colleagues’ 2014 paper with psychtoolbox in MATLAB [Wilson et al., 2014]. Participants played 192 games (in three blocks of 64 games) of the horizon task, which lasted around 60 minutes. Each gaming session consisted of either 5 or 10 trials, with the two-game lengths interleaved and counterbalanced to ensure an equal distribution of 96 games for each game length. Within each game, participants repeatedly made decisions between two options. These payouts were sampled from a Gaussian distribution with a consistent standard deviation of 8 points. The generative means of the underlying Gaussian differed for the two options and remained constant throughout each game. For each game, the mean of one option was fixed at either 40 or 60 points, while the mean of the other option was set relative to the first, with the difference between the two means being randomly sampled from 4, 8, 12, and 20 points.

Participants received verbal task instructions derived from the original task instructions outlined in the paper by [Wilson et al., 2014]. Overall, participants were directed to maximize their earned points and informed that one option consistently offered better outcomes on average.

Throughout each game, the choice and outcome history were displayed onscreen within each slot machine. Following the selection of a particular option, the reward for that trial was added to the corresponding slot machine, while the unchosen option’s space was filled with “X” (see Figure 1). The initial four trials of each game were designated as forced-choice trials, where participants were informed to select one of the options. These forced-choice trials established two information conditions: “unequal information” ([1 3]), requiring one option to be played once and the other three times, and “equal information” ([2 2]), with each option played twice. Following the forced-choice trials, participants made either one or six free choices.

**Fig. 1.**
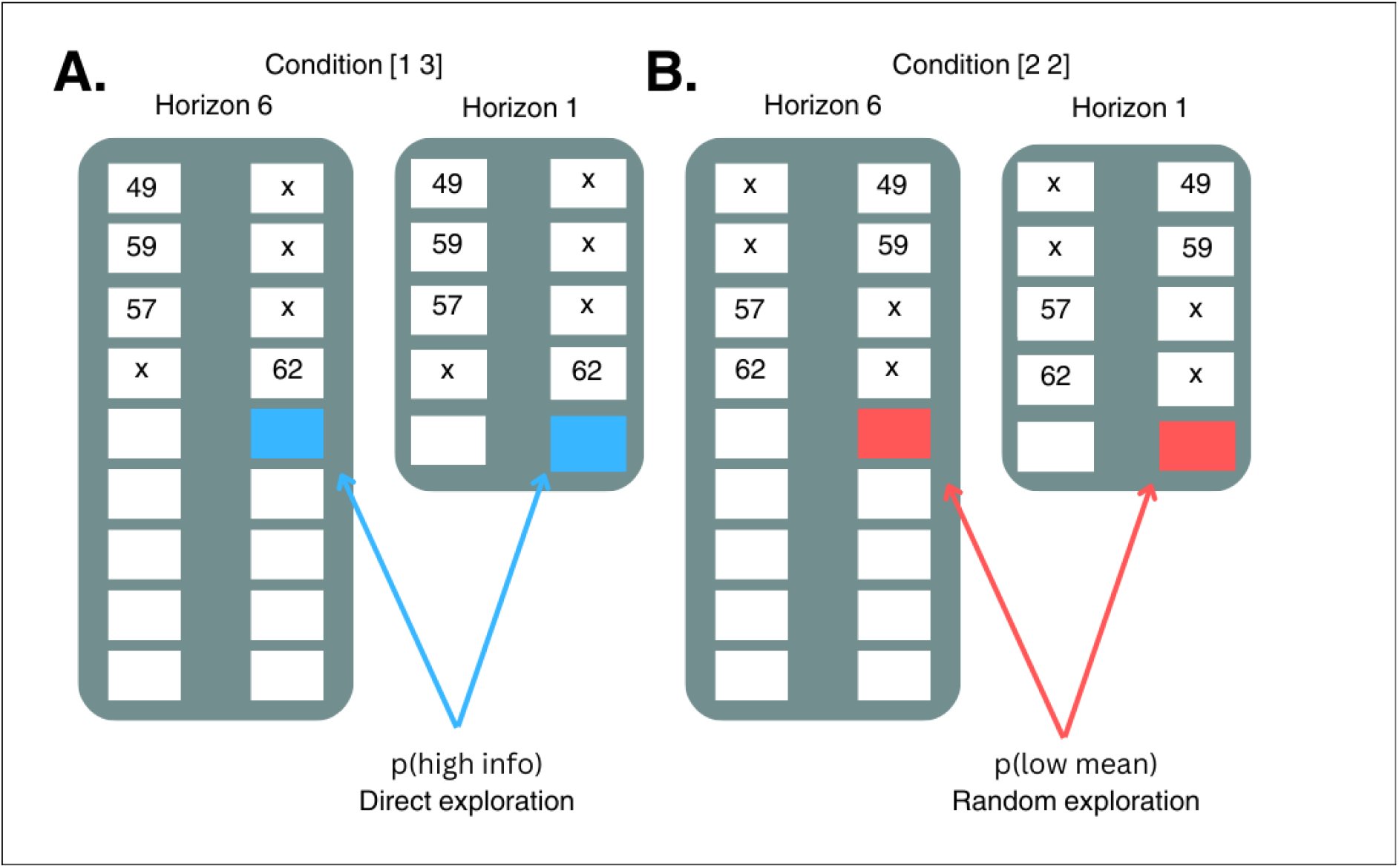
The horizon task. participants engage in decision-making between two armed bandits that offer probabilistic rewards. At the beginning, four ‘forced-choice’ trials provide partial information about the mean reward of each option. These trials are structured into two conditions with differing informational value: (A) In the unequal condition ([1 3]), participants observe one play from one option and three from another. Choosing the bandit with less information is considered a measure of information-seeking and direct exploration. (B) In the equal condition ([2 2]), both options provide the same amount of information, and selecting the option with the lower mean reward is considered a measure of random exploration.

**Fig. 2.**
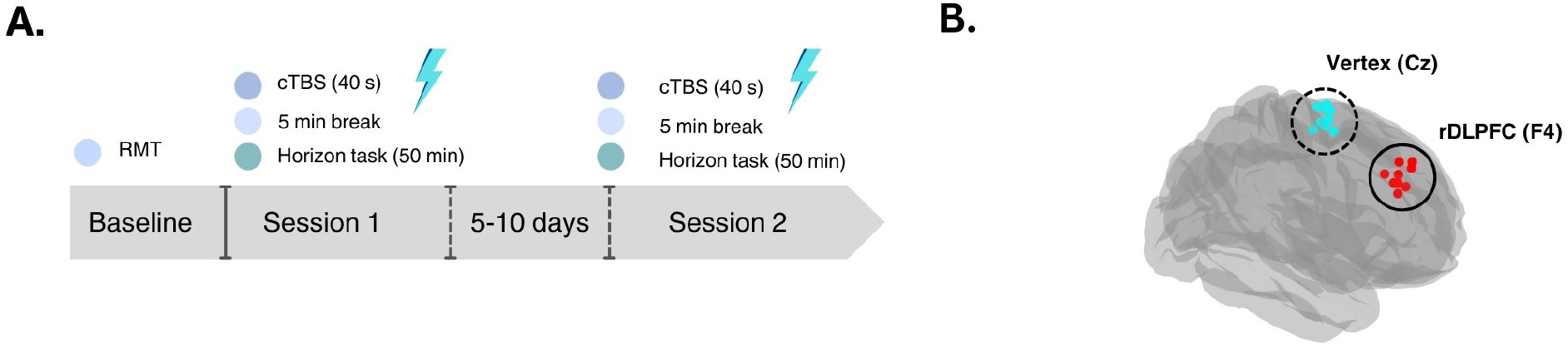
Experimental procedure. (A) experimental timeline. (B) stimulation conditions

The overall performance of participants throughout the experiment was evaluated to assess task performance. Any participant whose performance did not exceed the chance level at a threshold of p = .55 correct responses was excluded from the analysis.

### 2.5. Statistical Analysis

To quantify direct and random exploration, we employed established model-free and model-based approaches, focusing on the first free-choice trial to minimize confounding effects related to reward and information [Wilson et al., 2014, Zajkowski et al., 2017a]. Statistical analyses were conducted using MATLAB version 2022b.

In the model-free approach, directed exploration was assessed by calculating the probability of choosing the high-informative option, denoted as *p*_(high info)_, in the [1 3] condition. Random exploration was estimated as the probability of choosing the low-mean reward option, *p*_(low mean)_, in the [2 2] condition. Subsequently, a repeated measures ANOVA was performed with the horizon, TMS condition, order, and gender as factors, followed by post hoc analyses (paired t-tests) to examine the effect of TMS in different horizon conditions.

For the model-based approach, an established logistic choice rule model was applied to fit decisions made during the first free trial. This model posited that the value of each option (*a* or *b*) is influenced by three primary parameters: the average reward (R) associated with each option, the information bonus (I) provided by selecting an informative option, and spatial bias (s), reflecting potential biases in selecting options based on their location. The value of each option (*Q*_*a*_ or *Q*_*b*_) was expressed as:

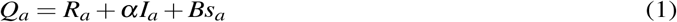

where *α* represents the information bonus, and B denotes spatial bias.

We further incorporated logistic noise with standard deviation *σ*_*d*_ to account for decision variability.

The probability of selecting option a over b was calculated as:

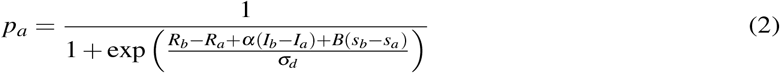

By fitting this model to our data, we estimated information bonus (*α*), spatial bias (B), and decision noise (*σ*_*d*_) in the different horizon and uncertainty conditions for each participant (2 parameters for information bonus and 4 parameters for decision noise and spatial bias). Then we perform a correlation between information bonus (*α*) and the model-free parameter *p*_(high info)_, as well as between decision noise (*σ*_*d*_) and *p*_(low mean)_. Furthermore, we assessed the proportion of correct responses across all trials, defined as selecting the option with the higher generative mean, and conducted paired t-tests to examine differences between the two TMS conditions.

## 3. Result

### 3.1. Model-free analysis

In the Horizon Task, the presence of two information conditions enables the measurement of both direct and random exploration in a model-free way. We conducted a repeated measures ANOVA with horizon, TMS condition, order, and gender as factors. In both direct and random exploration, we observed a significant main effect of the horizon condition (for directed exploration, *F*(1, 21) = 7.43, *p* = 0.001; for random exploration, *F*(1, 21) = 15.73, *p* < 0.001). This was attributed to the higher accuracy observed in the horizon 1 condition (*M* = 0.73, SD = 0.04), compared to horizon 6 (*M* = 0.66, *SD* = 0.02).

In random exploration, we also observed a significant effect of the stimulation condition (*F*(1, 21) = 9.73, *p* = 0.005). However, such an effect was not evident in direct exploration (*F*(1, 21) = 0.31, *p* = 0.58). Post hoc analyses revealed that the changes in random exploration were driven by changes in p(low mean) in both horizon 1 (paired t-test, *p* = 0.003) and horizon 6 (paired t-test, *p* = 0.046) conditions (Figure 3). Moreover, we found no gender or order effect in either random or direct exploration.

**Fig. 3.**
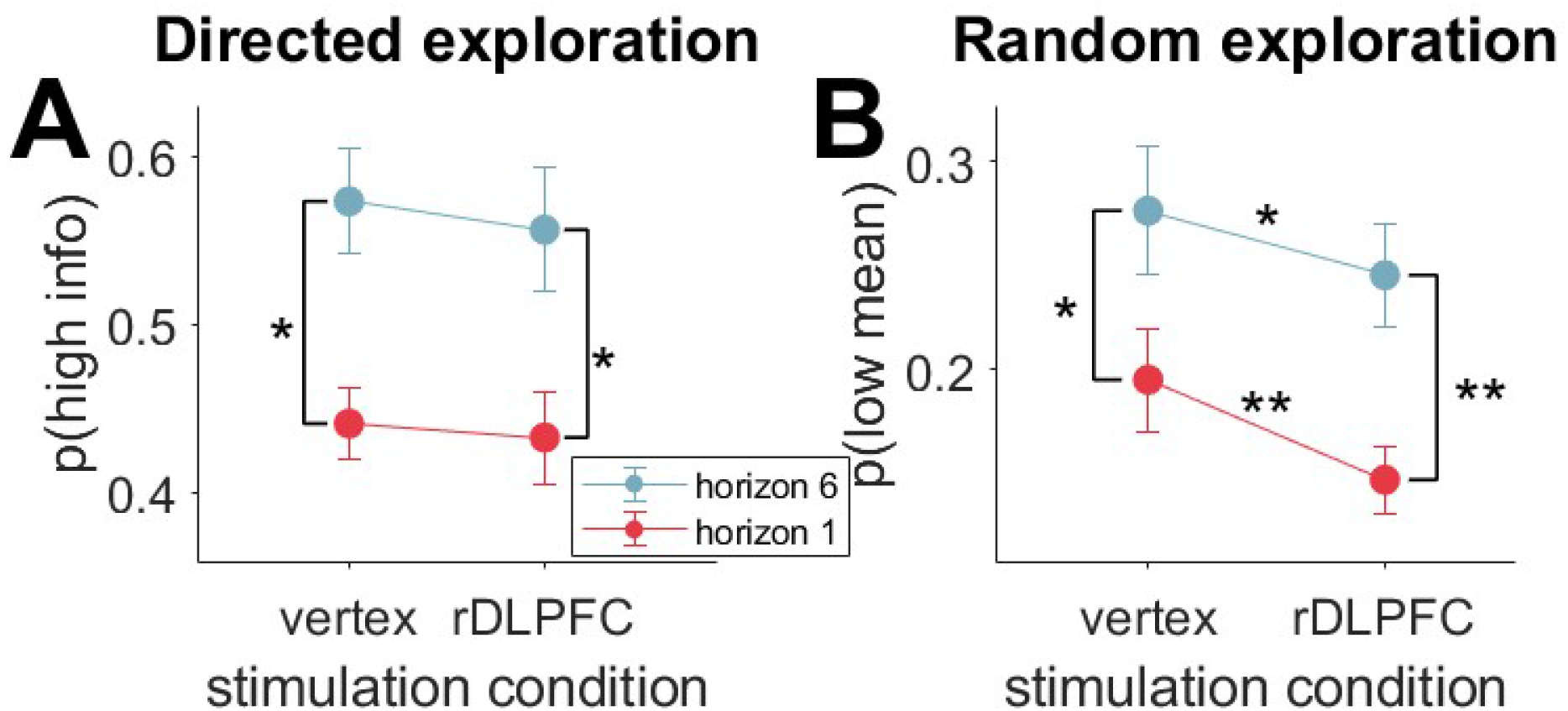
Model-Free Analysis of first Free-Choice Trials. (A) Stimulation of the right DLPFC via the cTBS protocol did not alter direct exploration across any horizon conditions. (B) Contrarily, random exploration significantly decreased following rDLPFC stimulation when compared to the control condition (vertex). In both scenarios, exploration increases with horizon. (* denotes *p* < 0.05, ** denotes *p* < 0.005; error bars represent standard error of the mean).

In our task, participants completed a total of 192 trials (three blocks of 64 trials), which typically took 50 minutes to complete. This duration might exceed the window of the TMS effect and could potentially induce fatigue. To address this concern, we conducted additional analyses on the probability of choosing the low-mean option (*p*(low mean)) and the probability of choosing the high-information option (*p*(high info)) within the first 64 and 128 trials.

The results revealed a significant reduction in random exploration even within the first 128 trials, both in horizon 1 (*p* = 0.01) and horizon 6 (*p* = 0.03) (see Figure 4). However, there were no significant changes in direct exploration observed in either the first 64 or 128 trials (see Supplementary Material 1), nor in random exploration within the first 64 trials. Additionally, an analysis of all free choices indicated no reaction time changes between stimulation conditions (Supplementary Material 4). Meanwhile, we found a considerable increase in the fraction of correct responses after the rDLPFC stimulation (see Figure 5). However, this change did not reach statistical significance (for horizon 1, p = 0.06).

**Fig. 4.**
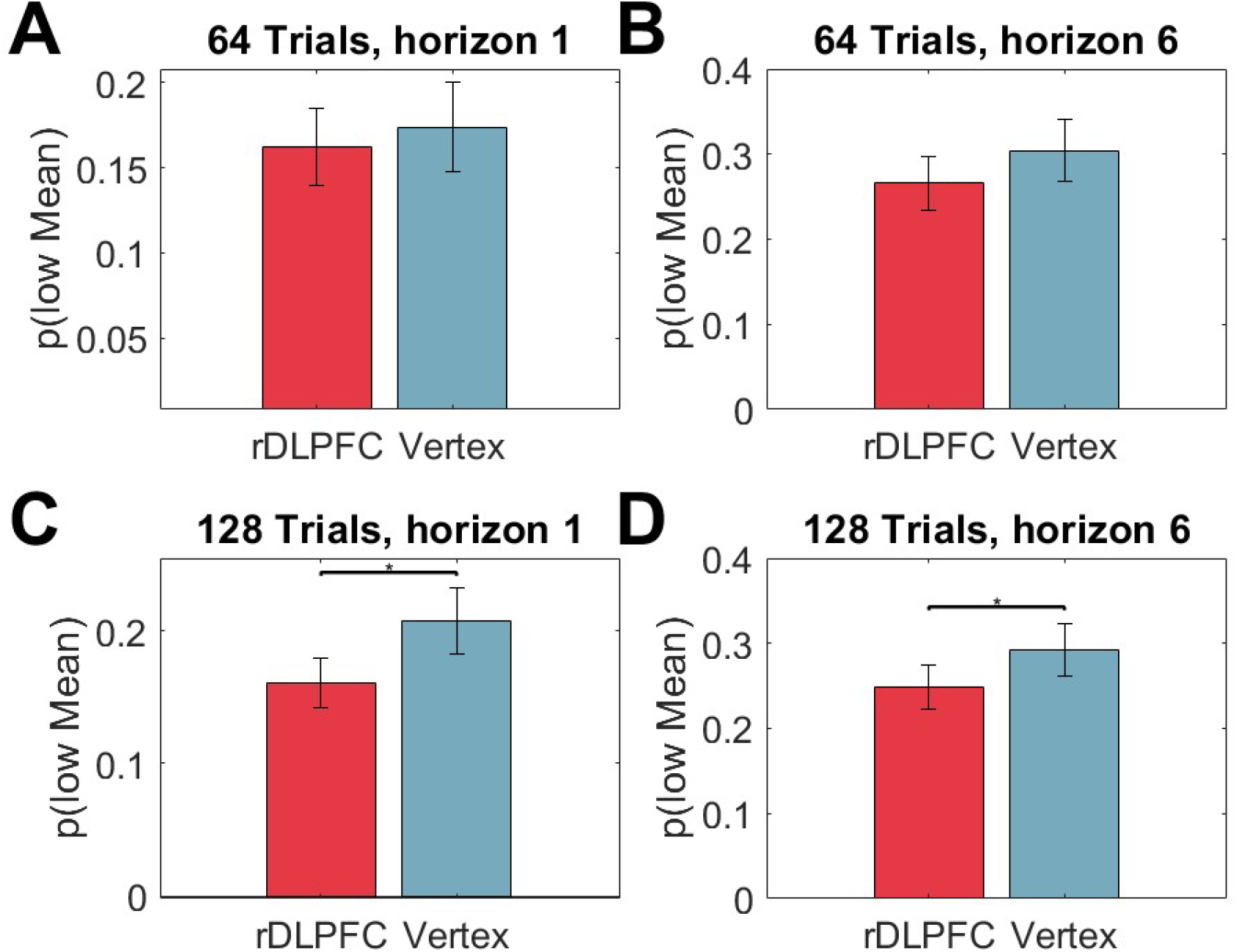
Model-Free Analysis of random exploration in first 64 and 128 trials. The first 64 trials showed no significant effect of rDLPFC stimulation on random exploration in either the horizon 1 (A) or horizon 6 (B) conditions. However, across the first 128 trials, rDLPFC stimulation led to a reduction in random exploration in both horizon conditions (C,D). (* denotes *p* < 0.05)

**Fig. 5.**
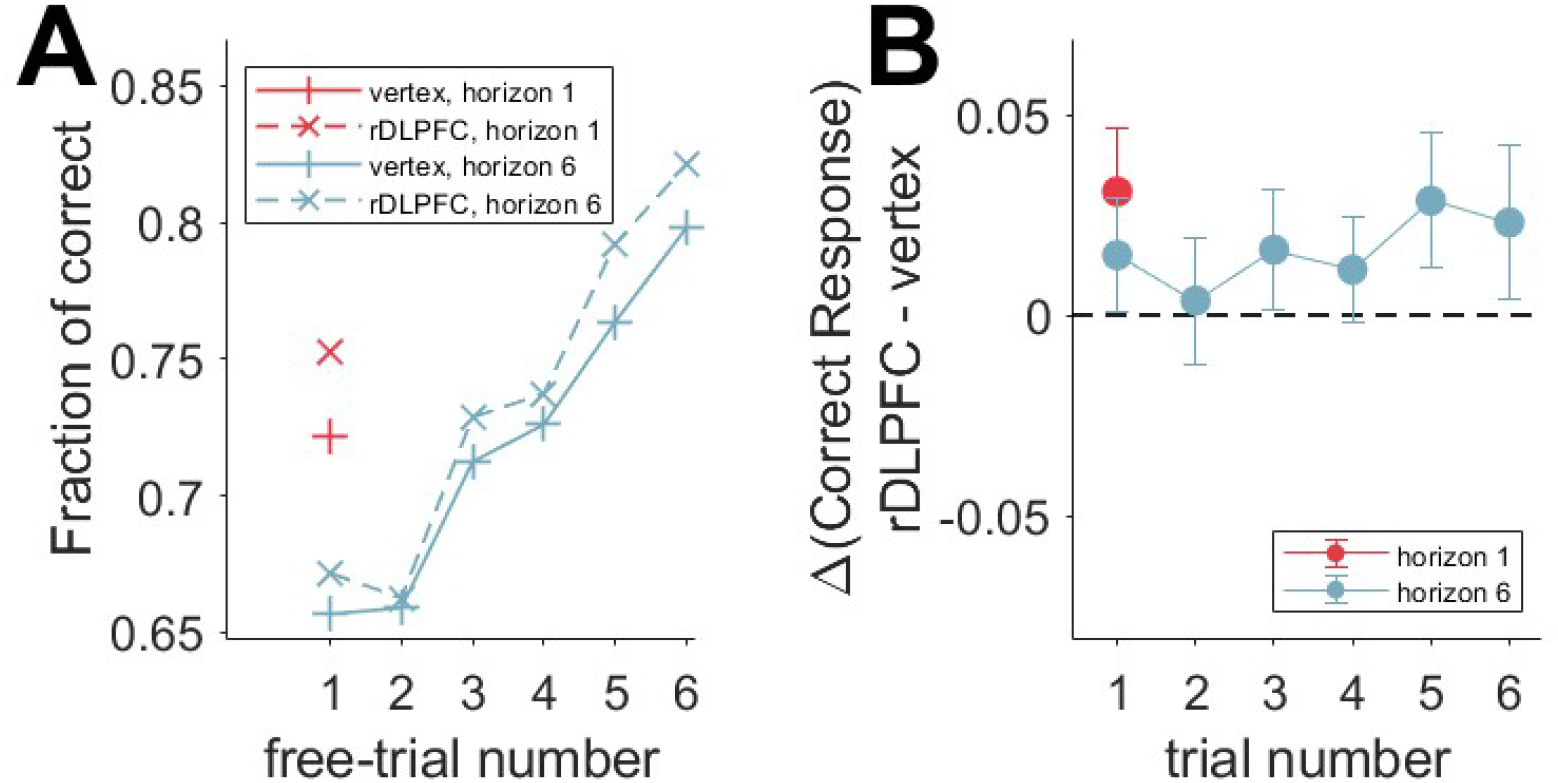
The fraction of correct responses of all free trials. (A) participants consistently performed above chance levels and exhibited improvement throughout the game. (B) stimulation of rDLPFC positively affected the accuracy of responses across all free trials, with a notable impact observed in the horizon 1 condition. However, these changes did not achieve statistical significance.

### 3.2. Model-Based analysis

We also employed a logistic choice rule model comprising three decision components: “information bonus,” “decision noise,” and “spatial bias.” These parameters were further categorized based on different horizons (1 or 6) and uncertainty conditions ([1 3] or [2 2]). Overall, our model consisted of 10 free parameters (four parameters for decision noise and spatial bias, and two parameters for information bonus). Meanwhile, the information bonus and decision noise correspond to direct and random exploration, respectively.

Results indicated a significant reduction only in decision noise for horizon 1, [2 2] condition (*p* = 0.042) and horizon 6, [2 2] condition (*p* = 0.025) (Figure 6). This effect was not observed for any other parameters in information bonus and spatial bias (see Supplementary Material 2 and 3).

**Fig. 6.**
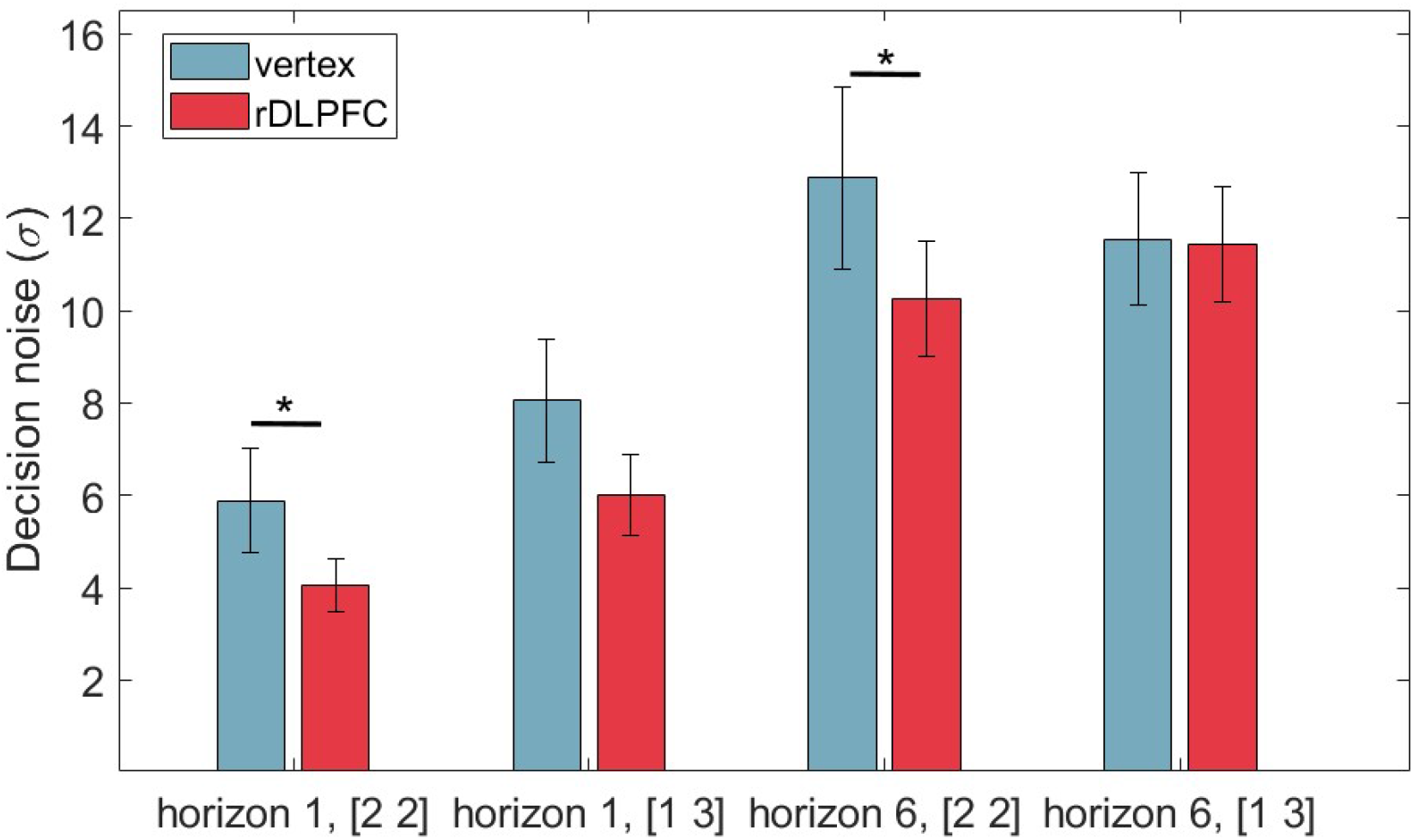
Model-Based Analysis of first Free-Choice Trials. Decision noise parameters for both the horizon 1 ([2 2]) condition and the horizon 6 ([2 2]) condition were significantly reduced following rDLPFC stimulation.

**Fig. 7.**
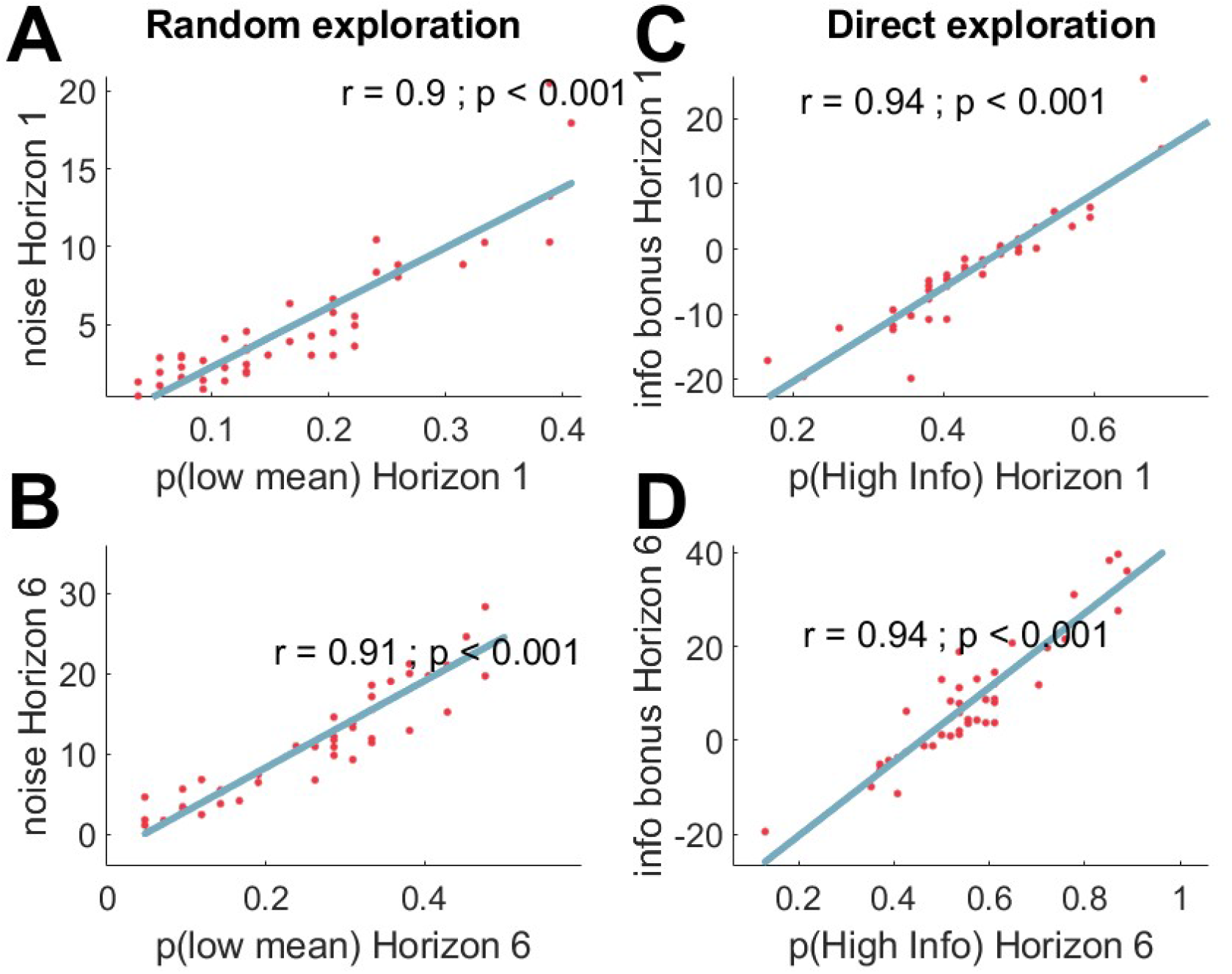
Correlations between model-based and model-free parameters. (A) decision noise in [2 2] condition (from model) and p(low mean) in horizon 1, (B) decision noise in [2 2] condition (from model) and p(low mean) in horizon 6, (C) information bonus (from model) and p(high info) in horizon 1, (D) information bonus (from model) and p(high info) in horizon 6

Finally, we conducted a correlation analysis between model-free and model-based results for both random and direct exploration. As expected, the results showed a high correlation between model-based and model-free analyses in both horizon conditions (Figure 5).

## 4. Discussion

In this study, we causally investigate the hypothesis that rDLPFC is involved in random and direct exploration. To address this hypothesis, we used continuous theta-burst transcranial magnetic stimulation (cTBS) over the right DLPFC to down-regulate its excitability and examine its effect on random and direct exploration while participants performed a horizon task. This task enables us to behaviorally dissociate these two types of explorative strategies, and remove reward-information confound by only analyzing the first free choice trial.

We found suppression of rDLPFC selectively reduced the random exploration, but not directed exploration, as evidenced by both model-free and model-based analysis. This study supports a previous fMRI study [Tomov et al., 2020] that showed rDLPFC in total uncertainty conditions, drives random exploration but not direct exploration and further supports the idea of different neural implementations for direct and random exploration.

The role of the rDLPFC in exploration is well-established in human and primate neuroimaging studies [Hogeveen et al., 2022b, Jahn et al., 2023, Tomov et al., 2020, Zhen et al., 2022], but our findings specifically highlight its involvement in random exploration. This raises two key questions: Firstly, why there were no observed changes in directed exploration? Secondly, what accounts for the decrease in random exploration?

Direct exploration is a bias toward more informative options. Previous studies that tried to disentangle direct and random exploration, did not find rDLPFC activation in direct exploration, instead they supported the role of dorsal ACC [Dezza et al., 2022] and rFPC [Badre et al., 2012, Tomov et al., 2020] specifically for direct exploration. These studies are aligned with our observation that showed rDLPFC stimulation didn’t change information-seeking behavior.

At the same time, direct exploration could also be linked to risk-taking behavior as choosing an option with high uncertainty could be interpreted as taking risks [Sadeghiyeh et al., 2020]. Previous noninvasive brain stimulation studies showed suppressing DLPFC, increases risk-taking behavior [Dantas et al., 2023, Knoch et al., 2006, Obeso et al., 2021], probably by affecting dopamine release in the striatum [Ko et al., 2008, Ott et al., 2011]. Complementary, dopamine modulation has been suggested to be involved in direct exploration [Chakroun et al., 2020]. At first glance, these studies seem at odds with our observation. However, the effect of cTBS in risk-taking behavior and increasing dopamine release seems to be limited to the left DLPFC (and not the right DLPFC) [Ko et al., 2008, Ott et al., 2011, Panidi et al., 2022]. A recent meta-analysis of fMRI studies related to exploration-exploitation-related tasks (e.g., armed bandit and foraging tasks) showed that rDLPFC activation is specific for exploration and not risk [Zhen et al., 2022]. In that sense, although risk-taking and exploration are two interdependent processes, their neural correlates are dissociable in some brain areas like the right DLPFC [Zhen et al., 2022].

Random exploration is defined as behavioral or neural variability, but the selective neural substrate for random exploration is less investigated [Wilson et al., 2021]. In an fMRI study conducted by Tomov et al. [2020] human participants performed a two-armed bandit task with risky (i.e., a fixed amount of reward) and safe (i.e., a variable amount of reward) options. They revealed that in total uncertainty conditions in which both bandits were risky or safe, bold signal in rDLPFC predicts trial variabilities in random exploration, but not direct exploration. In another study conducted by Jahn et al. [2023], monkeys performed a modified version of the horizon task in an MRI scanner. They saw when monkeys selected the low-value option, ACC/MCC and dlPFC had the greatest activity, which highlights the role of this network in decreasing the reliance on expected values and driving exploration to the low-value option [Jahn et al., 2023]. Moreover, this network interacts with the monoamine system including norepinephrine [Tervo et al., 2014], which previously supported its role in modulating noise in the decision process which leads to random exploration [Cremer et al., 2023, Dubois et al., 2021].

To interpret our results in light of these studies, it seems rDLPFC inhibition decreases random exploration by disrupting the DLPFC-ACC/MCC and locus coeruleus network which relaxes the effect of average reward between options and drives random exploration.

An alternative answer would be reducing the random exploration due to a decrease in the impulsivity level after rDLPFC stimulation. Previous studies showed that stimulating rDLPFC with cTBS protocol reduces impulsivity levels in delayed discounting tasks [Chakroun et al., 2020, Cho et al., 2010]. In a recent study, Dubois and Hauser [2022] showed a strong correlation between random exploration and general impulsivity in a large healthy sample size (N = 580) across multiple questionnaires. These studies could support the idea that cTBS over right DLPFC reduces random exploration by reducing impulsivity and enhancing self-control. At some point, this possibility also could be ruled out as a recent systematic review and meta-analysis of noninvasive brain stimulation over DLPFC, reveals a functional lateralization between right and left DLPFC which stimulation of left DLPFC and not rDLPFC, associated with reducing impulsivity, and enhancing self-control [Lin and Feng, 2024].

Lastly, random exploration could also be related to boredom [Wilson et al., 2021]. Thus, engaging in 192 trials might lead to disengagement from the task, and rDLPFC stimulation might improve the task performance by reducing fatigue [Soutschek and Tobler, 2020]. To test this hypothesis, we analyze the data in different blocks (the first 64 trials and the first 124 trials). Random exploration reduction is also evident in the first 124 trials in both horizon conditions (Figure 4). That’s why, it’s unlikely that these changes happen due to fatigue prevention.

These results could also have potential clinical implications, given the known associations between exploration-exploitation imbalances and various psychopathologies such as schizophrenia [Cathomas et al., 2021, Speers and Bilkey, 2023, Strauss et al., 2011], depression [Blanco et al., 2013, Smith et al., 2022], and addiction [Addicott et al., 2017]. Specifically, the overuse of random exploration observed in schizophrenia patients, which correlates with poorer decision-making performance [Cathomas et al., 2021, Speers and Bilkey, 2023], suggests a targeted therapeutic opportunity. Future research could explore interventions that can specifically modulate cortical excitability of the rDLPFC to possibly modify random exploration imbalance.

It should be mentioned that this study has a notable limitation. Our experimental design lacks spatial resolution as we don’t use MRI coordinates for stimulating rDLPFC. In an ideal design, one could use the MRI coordinates of previous neuroimaging studies, to specify which part of the right DLPFC is exactly stimulated. Future studies utilizing fMRI and TMS are needed to replicate results and unravel the possible network interaction supporting random exploration.

In summary, our study for the first time provides causal evidence for the involvement of the right DLPFC in random exploration. This result is aligned with previous neuroimaging studies and further supports the dissociable neural implementation for direct and random exploration.

---

**Table 1.** Summary of Participant Characteristics.

| Characteristic | Mean | Standard Deviation |
| --- | --- | --- |
| Number of Participants | 25 (11 female) | - |
| Age (years) | 23.84 | 2.76 |
| RMT | 53.52 | 5.23 |
| DASS-21-total | 9.2 | 5.57 |

## Notes

https://github.com/ArminTi/A-causal-role-of-rDLPFC-in-random-exploration

### Competing Interest Statement

The authors have declared no competing interest.

