## Supplementary material for "A Causal Role of the Right Dorsolateral Prefrontal Cortex in Random Exploration"

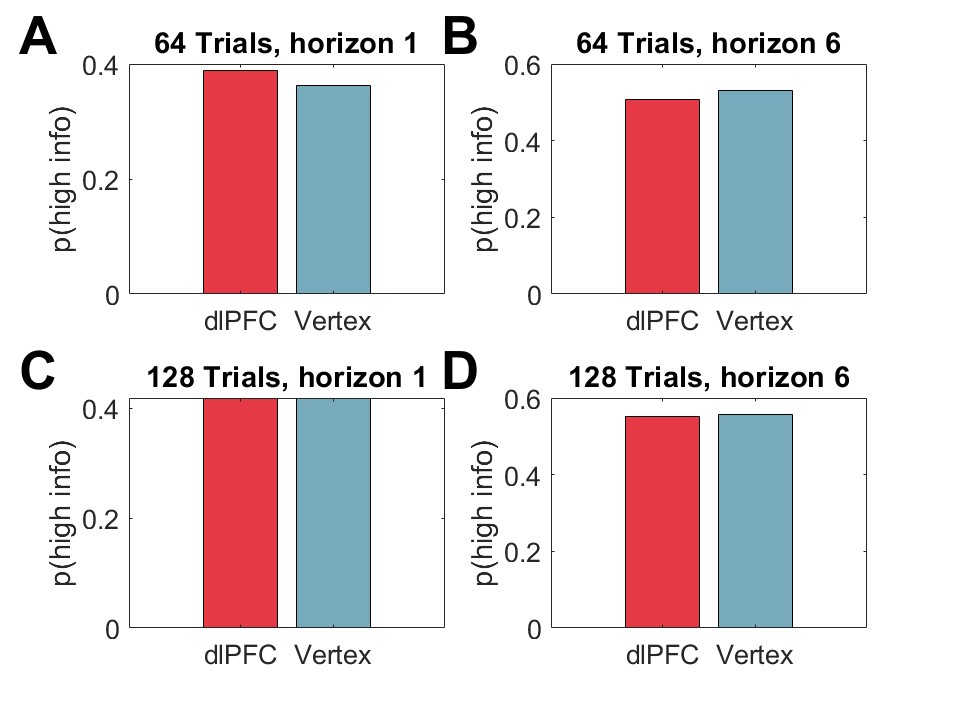

Figure S1: Model-free analysis of direct exploration in the first 64 and 128 trials. The first 64 and 128 trials showed no significant effect of rDLPFC stimulation on direct exploration in horizon 1 (p = 0.51 (A), p = 0.96 (C)) or horizon 6 (p = 0.58 (B), p = 0.83 (D)) conditions.
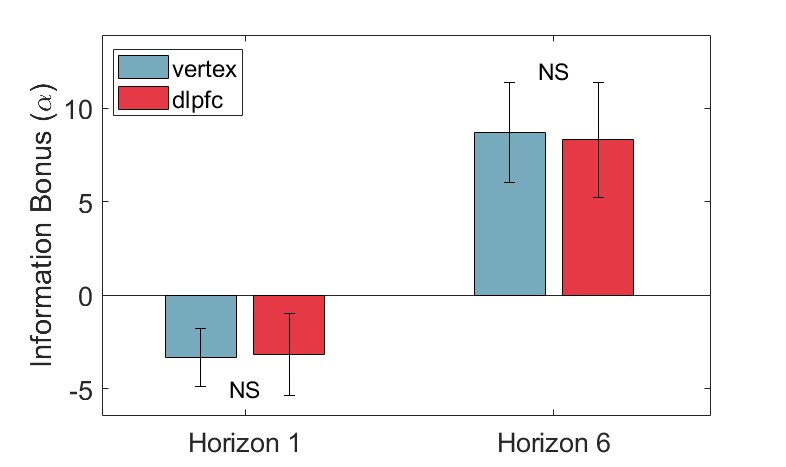

Figure S2: Model-Based Analysis of First Free-Choice Trials. Information Bonus parameters for
both the horizon 1 condition (p = 0.95) and the horizon 6 condition (p = 0.85) were not significantly different after rDLPFC stimulation.

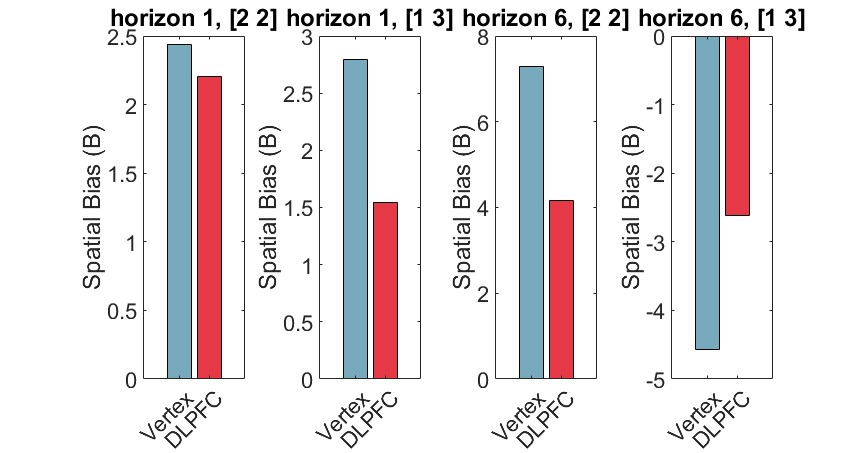

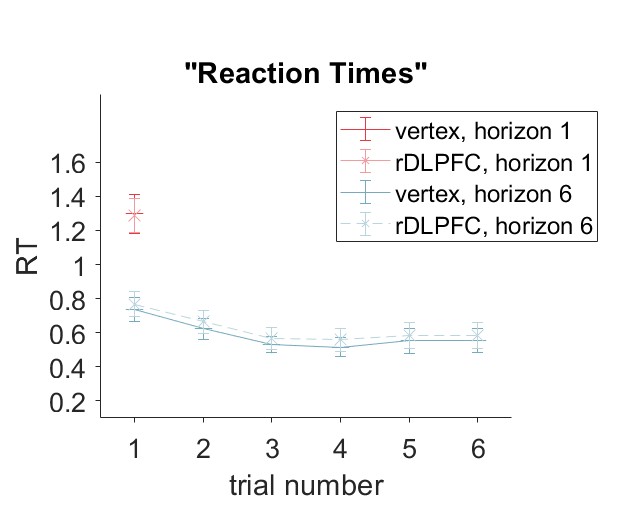
Figure S3: Model-Based Analysis of First Free-Choice Trials. Spatial bias parameters for
both the horizon 1 condition (for [2 2], p = 0.83, for [1 3], p = 0.56) and the horizon 6 conditions (for [2 2], p = 0.07, for [1 3], p = 0.42) were not significantly different after rDLPFC stimulation.

Figure S4: Reaction times in all free-choice trials. There was no significant change in reaction times after stimulation.
